# Altering the solubility of the antibiotic candidate Nisin – a computational study

**DOI:** 10.1101/2020.07.16.206821

**Authors:** Preeti Pandey, Ulrich H.E. Hansmann, Feng Wang

## Abstract

The growing bacterial resistance to available antibiotics makes it necessary to look for new drug candidates. An example is a lanthionine-containing nisin, which has a broad spectrum of antimicrobial activity. While nisin is widely utilized as a food preservative, its poor solubility and low stability at physiological pH hinder its use as an antibiotic. As the solubility of nisin is controlled by the residues of the hinge region, we have performed molecular dynamics simulations of various mutants and studied their effects on nisin’s solubility. These simulations are complicated by the presence of two uncommon residues (dehydroalanine and dehydrobutyrine) in the peptide. The primary goal of the present study is to derive rules for designing new mutants that will be more soluble at physiological pH and, therefore, may serve as a basis for the future antibiotic design. Another aim of our study is to evaluate whether existing force fields can model the solubility of these amino acids accurately, in order to motivate further developments of force fields to account for solubility information.

## Introduction

The growing resistance to antibiotics, caused by its injudicious use and over-utilization in humans and animals, has become a threat to the global health care system and the safety of the food supply. New drugs are desperately needed to control the increasing spread of infectious diseases in humans and farm animals. Promising candidates are antimicrobial peptides (AMPs) produced by bacteria that target peptidoglycans in bacterial cell walls as they show little cross-resistance.^1,2^ Such bacteriocins are, in general, only effective against closely related strains. An exception is nisin, produced by the gram-positive bacterium *Lactococcus lactis*, which exhibits a broad-spectrum of antibacterial activity due to its stability at higher temperature, tolerance to low pH, and dual mode of action. When combined with high temperature and chelating agents, it is effective even against gram-negative bacteria, which makes nisin one of the most utilized food-preservatives in the world.^3^

Nisin belongs to the class of lantibiotics, characterized by uncommon residues such as dehydrated and lanthionine residues. The amphipathic 34-residue long peptide carries an overall positive charge, and its sequence of amino acids contains two unusual amino-acids, dehydroalanine (Dha) and dehydrobutyrine (Dhb), and five thio-ether rings (one lanthionine and four threo-β-methyl lanthionine), see Figure 1. The antimicrobial activity of nisin relies on two modes: “Lipid II trapping” and “Pore formation”. The first mode is due to the N-terminal residues (1-12) binding to the pyrophosphate moiety of lipid II,^4^ which carries the peptidoglycan unit, thus competing with the biosynthesis of the cell wall. At the same time, the C-terminus of nisin enables the formation of pores in the cell membrane, thus causing cell leakage resulting in cell death.^5^

**Figure 1.**
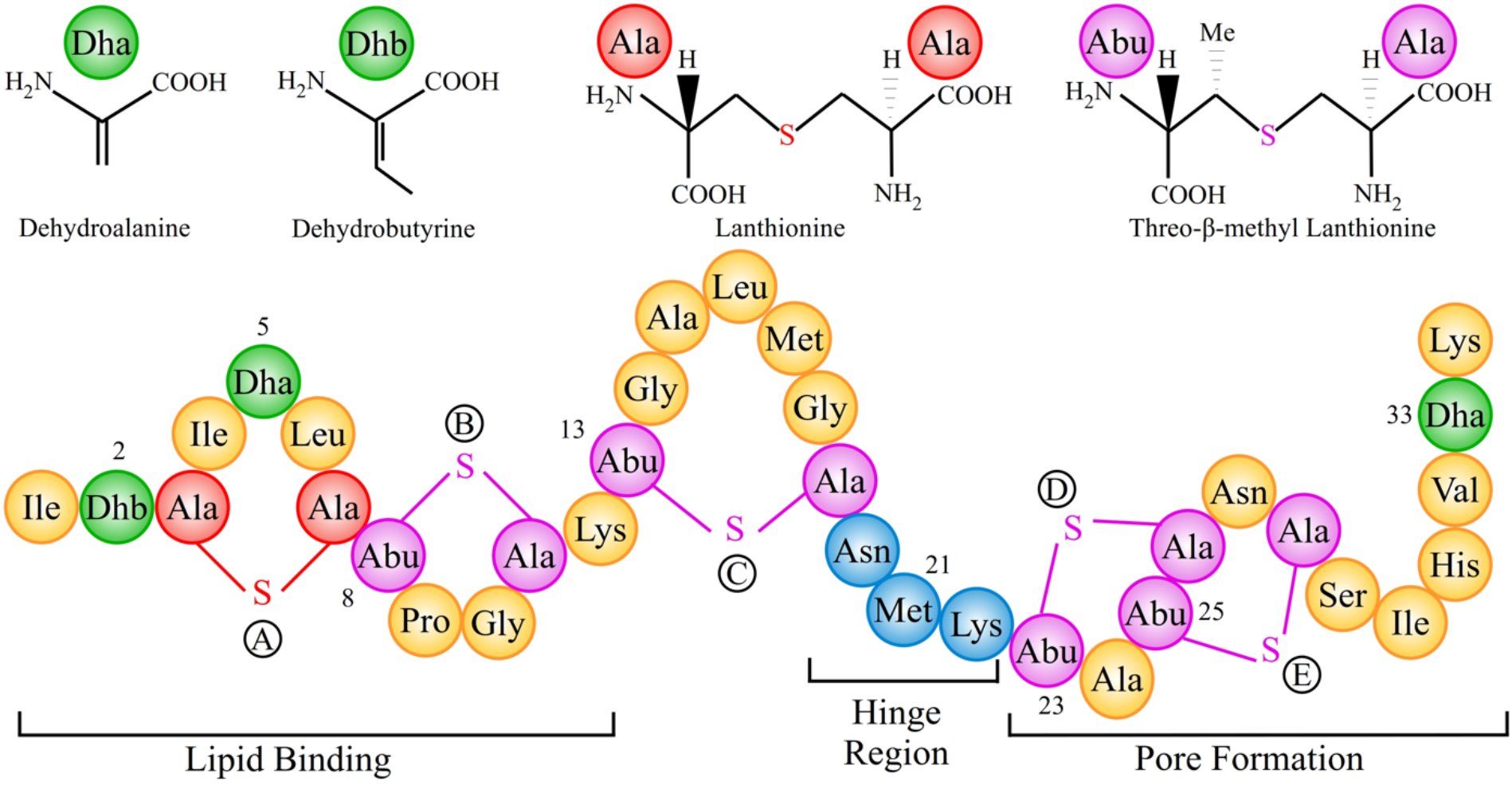
Bottom: primary structure of Nisin, highlighting the regions involved in distinct aspects of nisin’s antimicrobial activity. Top: Chemical formula of dehydroalanine (Dha), dehydrobutyrine (Dhb), and lanthionine rings.

Despite its antimicrobial activity and non-toxicity, the use of nisin as a food preservative is restricted by its poor solubility, low stability at physiological pH, and high temperature.^6,7^ These factors restrict even more the possible employment as an antibiotic. The solution structure of the nisin-3LII complex (nisin in complex with lipid II) and homology modeling of several related lantibiotics indicates that mainchain interaction between nisin and lipid II dominate. The lesser importance of sidechain interactions, therefore, provides an opportunity to design mutants with improved solubility that do not interfere with the lipid II binding of the peptide. For instance, Rollema *et. al*^8^ have reported two mutants, N27K, and H31K, that have similar activity as wild-type nisin but higher solubility at physiological pH (pH: 7). In a similar vein, Yuan *et. al*,^9^ altered various residues in the hinge region of nisin by site-directed mutagenesis and showed that the hinge region is essential for the conformational flexibility necessary for disrupting the bacterial membranes as well as controls the solubility of the peptide. Two mutants, N20K and M21K, have three-fold, and five-fold higher solubility than the wild type at a pH of 8; however, the double mutant N20K-M21K has lower solubility. They also reported other mutants such as N20Q and M21G that are at higher temperatures and neutral or alkaline pH considerably more stable than the wild-type. Nevertheless, despite these reports of mutants with improved solubility and higher stability at neutral pH, there exists no procedure for a rational design of mutants bearing all these properties, and specifically, the cause for the unexpected solubility of the reported mutants remains unclear.

A computational investigation of these questions is not only less costly and faster than the mutagenesis experiments but also gives insight into the energetics responsible for the observed solubilities. One problem that hampers the computational investigation of nisin is the presence of dehydroamino acids, and the thio-ether bridges are not parametrized in standard force fields such as CHARMM used in this study. Two sets of CHARMM compatible parameters have been proposed for the dehydroamino acids and the thio-ether bridges by Turpin *et. al*^10^ and de Miguel *et*. *al*;^11^ however, solubility predictions based on these models were never checked and validated. Hence, we investigate in the present work first the effect of force field on the solubility of nisin, using all-atom molecular dynamics simulations and solvation free energy calculations that rely on Poisson– Boltzmann and surface area continuum solvation (PBSA) and Generalized Born and surface area continuum solvation (GBSA) methods. In a second step, we propose three new mutants of nisin with improved solubility (N20K^+^-M21K^+^ (KK-PP), N20R-K22R and N20R), and introduce rules for predicting the solubility of nisin mutants as needed for future antibiotic design.

## Materials & Methods

### Design of mutants and simulation set

To understand the factors that alter the solubility of nisin and to study the extent by which the force field parametrization of the uncommon dehydroamino acids determines structural properties and solubility of nisin, we have carried out multiple molecular dynamics (MD) simulations of both wild type and suitable mutants. The mutations that we have studied are located in the hinge region of the peptide (residues 20-22) and are listed in Table 1. Note that lysine, arginine, and histidine in the mutants are protonated (*i.e.*, positively charged), while glutamic acid is deprotonated (*i.e.*, negatively charged). For clarity, the protonation states of Lysines are marked explicitly.

**Table 1.** List of mutants of Nisin.

|  | **Mutants** |
| --- | --- |
| **Positively Charged** | N20K<sup>+</sup> (denoted as N20K), M21K<sup>+</sup> (denoted as M21K), N20K-M21K (denoted as KK-NP), N20K<sup>+</sup>-M21K<sup>+</sup> (denoted as KK-PP), N20R, K22R, N20R-K22R, N20H, M21H, K22H |
| **Negatively Charged** | N20E, M21E, and K22E |
| **Uncharged Polar** | N20Q |
| **Hydrophobic** | N20V, N20F, M21G, K22G |
| **Other** | N20F-M21L-K22Q (denoted as FLQ) |

The initial coordinates of wild-type nisin required for the molecular dynamics simulations were obtained from the nisin-lipid II complex as resolved by NMR and deposited in the Protein Data Bank (PDB) (PDB Id: 1WCO).^4^ The three models with the lowest energy were selected from the deposited 20 solution NMR structures. After deletion of the model lipid II, mutants were derived for each of the three nisin models by altering the corresponding residue. In this way, we minimize the possibility that our results depend on the specifics of the selected NMR model and take into account the inherent flexibility of the peptide. Containing five thio-ether bridged rings and two uncommon amino acids, dehydroalanine (Dha) and dehydrobutyrine (Dhb), nisin does not attain regular secondary structural elements like α-helices and β-sheets. Computationally, this adds the difficulty that carefully designed parameters for the uncommon amino acids have to be added to the existing protein force fields in order to ensure reliable simulations. In the case of the CHARMM force field, two sets of parameters have been proposed for these unusual amino acids (Dha and Dhb) and the thio-ether bridges, one by Turpin *et. al*^10^ and one by de Miguel *et. al*.^11^ The Turpin parameters were fitted for the E-isomer of Dhb by using scaled Hartree-Fock for partial charges and MP2/6-31G*//MP2/cc-pVTZ for torsional and other intramolecular parameters; and validated on hydrated nisin to adequately reproduce experimental NMR data and hydrogen bonds information for nisin. The de Miguel parameters were fitted using the Z-isomer commonly observed in AMPs, following the same procedure as done by Turpin to be compatible with the remaining CHARMM parameters. The parameters were validated by simulating Lantibiotics such as Lchα and Lchβ but they were not tested specifically for nisin. Especially, the suitability of the parameters for estimating solubility was not tested during the development of either of the parameter sets. In order to compare the suitability of the two sets, we, therefore, carried out two sets of molecular dynamics simulations, both using the CHARMM36^12^ force field for standard amino acids but differing in the parameters for dehydroamino acids and the thio-ether bridges.

We use for our molecular dynamics simulations the software package GROMACS-2018.1.^13,14^ Each peptide was put into the center of a cubic box with a minimum peptide-box distance of 15 Å; then the box was subsequently filled with TIP3P water^15^ and 0.15 M NaCl, including neutralizing counter-ions. Because of periodic boundary conditions, electrostatic interactions are evaluated by particle-Ewald summation,^16,17^ and a cut-off of 12 Å was used to calculate vdW-interactions. The resulting systems were energy-minimized by steepest descent, followed by equilibration of 500 ps at constant volume and subsequent 500 ps at constant pressure. Temperature and pressure was regulated by a Parrinello-Danadio-Bussi thermostat^18^ and Parrinello-Rahman barostat^19^ and set to 300 K and 1 bar, respectively. The integration step was 2 fs. Production runs were performed for 200 ns, and the coordinates were saved every 1 ps for subsequent analysis. This resulted in a total of 24 μs data.

### Solvation free energy estimation

The solubility of the nisin and mutants is described by the solvation free energy of the peptides. Two popular and computationally efficient continuum solvation models, Poisson Boltzmann Surface Area (PBSA) and Generalized Born Surface Area (GBSA) calculations,^20,21^ were employed by us for estimating the solvation free energy of the peptides. Both methods work by taking snapshots from a molecular dynamics (MD) trajectory and replacing the explicit water by a dielectric continuum. In PBSA, the polar part of the solvation energy, *i.e.*, the electrostatic component is evaluated by solving the Poisson equation (if there is no salt) or Poisson-Boltzmann equation (if salt is present in the system). Here, the Poisson equation models the variation of the electrostatic potential in a medium with a uniform dielectric constant ε, and the Boltzmann distribution governs the ion distribution in the system. GBSA differs from PBSA in that the electrostatic contribution is estimated using the Generalized Born approximation. In both methods, the non-polar component of the solvation free energy is approximated by a surface-area based approach. For details of the two methods, see, for instance, Ref. 22. Both approaches are implemented in the MMPBSA.py^23^ script as available in AmberTools18.^24^ We used this script for our analysis, extracting evenly spaced snapshots from the last 50 ns of the production runs. A salt concentration of 0.15 M is used in all calculations, and the solute dielectric constant, ε_in_, is set to 4 in PBSA calculations.

### Statistical tests

Both assessing the suitability of the two force fields variants by comparing computational and experimental results and comparing the solubility of the nisin and its mutant forms requires careful choice of an appropriate statistical test. While the t-test is good for comparing the difference between means of two groups, it can result in Type-I error (occurs when H_0_ is statistically rejected even though it is true), also known as “family-wise error”, when performing multiple pairwise comparisons.^25^ Hence, a t-test was used only for comparison of the two force field variants, while in contrast, the multiple comparison tukey’s test is used to differentiate the solubility of the nisin and its mutant forms. This is because tukey’s test is more robust and precise since the variance is estimated from the whole data set as a pooled estimate. In addition, tukey’s test adjusts the p-values for multiple comparisons, and, therefore, controls the family-wise error rate.

## Result & Discussion

### Effect of force field on structural properties and solubility of nisin

The activity of antimicrobial peptides is determined by their secondary structure, charge, hydrophobicity, and solubility. While most have random disordered configurations in an aqueous environment, they fold into an α-helix or β-sheet when interacting with membranes. Given the importance of secondary structure and solubility for the functions of antimicrobial peptides, it is important for the computational studies to employ force fields that do not introduce a bias into either structural properties or solubility of the peptides. This is especially a concern for nisin as it has dehydroamino acids and the thio-ether bridges, which are not parametrized in standard implementations of most force fields such as CHARMM36 used by us. Hence, we have to rely on modifications such as the ones proposed by Turpin *et. al*^10^ and de Miguel *et. al*.^11^ Hence, we first evaluate the suitability and limitations of these two parametrizations by probing their effect on structural properties and solubility of nisin.

The NMR model of nisin (with 20 downloaded solution structures in the PDB under identifier 1WCO) has a stable N-terminus and a flexible C-terminal tail.^4^ The N-terminus was stabilized into the cage-like structure by binding to the lipid II model. Since the lipid II was deleted in our simulation, we anticipate nisin to exhibit little canonical secondary structure. This can be seen in Figure 2, where we compare the root-mean-square-fluctuations (RMSF) of each residue using both the parameter sets (Figure 2). Although both parameters sets lead to similar trends, the fluctuations of the residues are larger in the simulations relying on the parameters of de Miguel *et. al* than in the ones that used the set proposed by Turpin *et. al*. Similarly, Figure S1 shows that the distribution of Phi-psi (ϕ/ψ) backbone dihedral angles are broader when using de Miguel *et. al* parameters for the dehydroamino acids while when using Turpin *et. al* parameters the distribution of dihedral angles is more narrow indicating sampling of a more restricted set of backbone conformations. This is consistent with the observation that the structural ensembles obtained using Turpin *et. al* parameters resemble more closely the ensemble of NMR structures than the ensembles obtained from simulations relying on the parameters of de Miguel *et. al*. This is not surprising considering that the de Miguel parameters had not been validated specifically for nisin simulations, but it does not necessarily indicate the de Miguel parameters to be of lower accuracy since increased flexibility is anticipated upon removal of the lipid II. While the NMR results suggest a stable small 3_10_ helix conformation at the N-terminus that might be essential for binding of nisin to lipid II, we did not observe such a 3_10_ helix in our simulations with either parameter set. Instead, we find at the N-terminus for both parameters set only flexible turns and random coil conformations. This result is consistent with circular dichroism measurements, which indicate that nisin typically assumes a random coil configuration in aqueous environment.^26^ This suggests that nisin adopts a 3_10_ helix only when bound to lipid II, or that this secondary structure results from the combined effect of lipid II binding and the use of DMSO (Dimethyl Sulfoxide) during structure determination. Hence, we believe that the absence of the 3_10_ helix in our simulation does not point to shortcomings of the two parameter sets.

**Figure 2.**
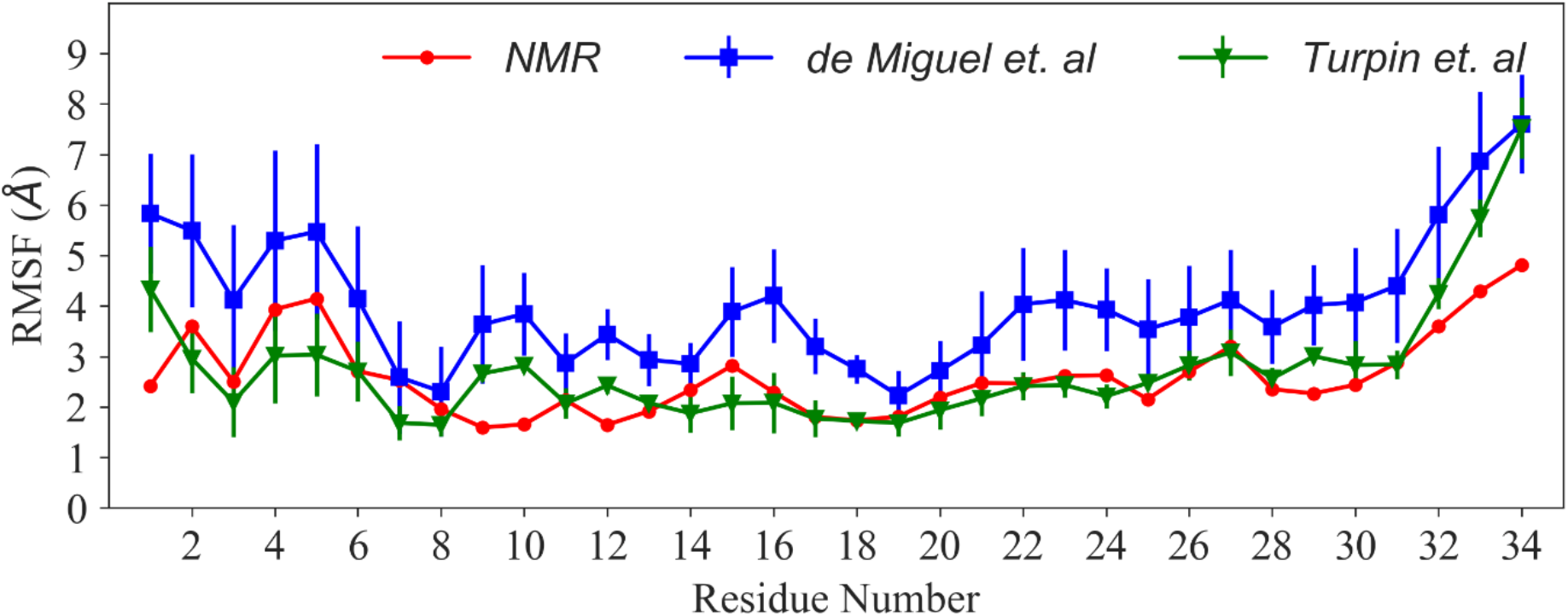
Root-mean-square fluctuation (RMSF) of Cα-atoms of Nisin.

Comparing solubility measurements with the calculated solvation free energy of nisin and its mutants allows one to probe the performance of these force fields in reproducing experimental trends. As described in the method section, we estimate the solvation free energies by PBSA and GBSA approximations. Using Welch’s t*-*test for comparing the numerical results obtained from simulations with the two parameter sets, we find significant differences (p-value < 0.001) between the two parameter sets for the solvation free energies (ΔG_sol_) of the wild-type and mutants. The only exceptions are the KK-PP mutant (where the GBSA values agree) and the N20R mutant (where both GBSA and PBSA values agree), see also Table 2 and Table S1. Note also that although the differences between the two parameter sets are small in the averages, there are substantial differences in the standard deviations, which with only a few exceptions, are always larger with de Miguel *et. al* parameters. Table 2 also shows that solvation free energy estimates obtained by GBSA and PBSA agree with each other, once one accounts for the choice of the solute dielectric constant. With an ε_in_ of 4 in PBSA, ΔG_sol_ values (and the corresponding standard deviations) computed with PBSA is four times smaller than the corresponding values obtained using GBSA. While the GBSA and PBSA in general lead to similar results, the case of the KK-PP mutant, where the GBSA values, but not the PBSA values, agree for both force fields, may point to the possibility that GBSA overestimates the polar component of the solvation energy, a problem that can be easily addressed in PBSA by using a more realistic internal dielectric constant (**ε_in_**). Hence, despite GBSA being the less costly approach, we focus in the following analysis on our PBSA data.

**Table 2.**
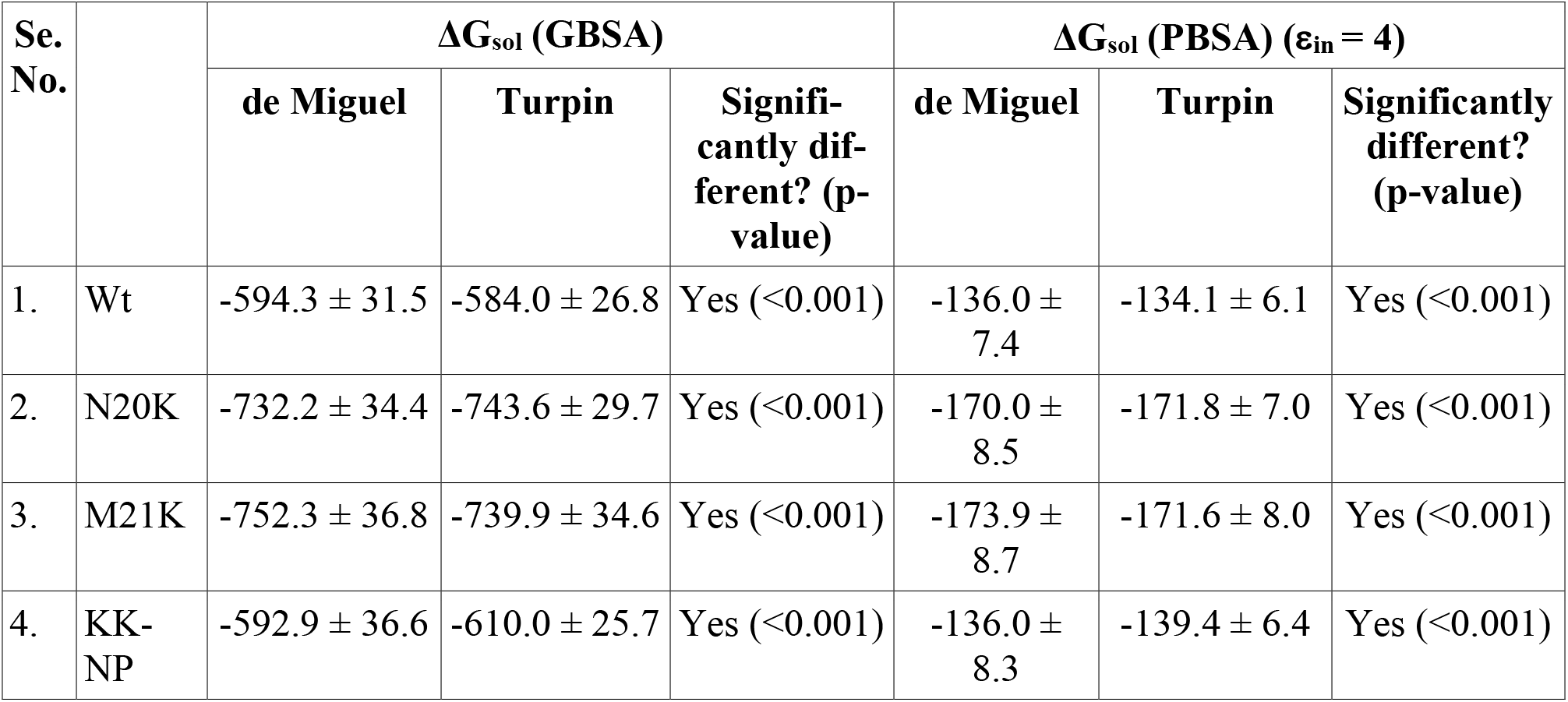
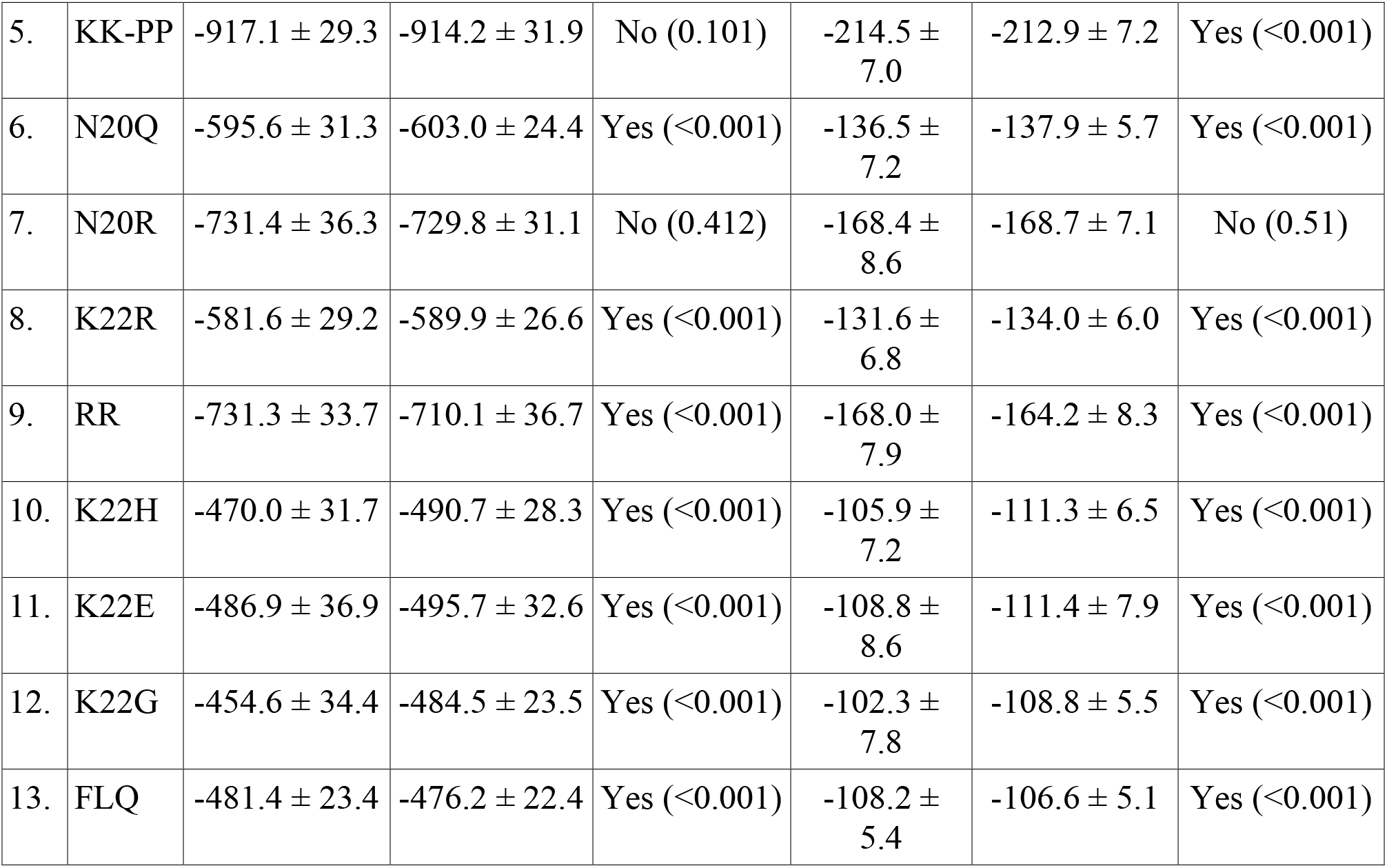
Calculated p-values of statistical test (Welch’s t-test) for solvation free energy (GBSA and PBSA) of Nisin and its mutant forms using parameters of de Miguel *et. al* and Turpin *et. al.*

| Se. No. | | $\Delta G_{sol}$ (GBSA) | | | $\Delta G_{sol}$ (PBSA) ($\epsilon_{in} = 4$) | | |
| --- | --- | --- | --- | --- | --- | --- | --- |
|  |  | de Miguel | Turpin | Significantly different? (p-value) | de Miguel | Turpin | Significantly different? (p-value) |
| 1. | Wt | -594.3 ± 31.5 | -584.0 ± 26.8 | Yes (<0.001) | -136.0 ± 7.4 | -134.1 ± 6.1 | Yes (<0.001) |
| 2. | N20K | -732.2 ± 34.4 | -743.6 ± 29.7 | Yes (<0.001) | -170.0 ± 8.5 | -171.8 ± 7.0 | Yes (<0.001) |
| 3. | M21K | -752.3 ± 36.8 | -739.9 ± 34.6 | Yes (<0.001) | -173.9 ± 8.7 | -171.6 ± 8.0 | Yes (<0.001) |
| 4. | KK-NP | -592.9 ± 36.6 | -610.0 ± 25.7 | Yes (<0.001) | -136.0 ± 8.3 | -139.4 ± 6.4 | Yes (<0.001) |
| 5. | KK-PP | $-917.1 \pm 29.3$ | $-914.2 \pm 31.9$ | No (0.101) | $-214.5 \pm 7.0$ | $-212.9 \pm 7.2$ | Yes (<0.001) |
| 6. | N20Q | $-595.6 \pm 31.3$ | $-603.0 \pm 24.4$ | Yes (<0.001) | $-136.5 \pm 7.2$ | $-137.9 \pm 5.7$ | Yes (<0.001) |
| 7. | N20R | $-731.4 \pm 36.3$ | $-729.8 \pm 31.1$ | No (0.412) | $-168.4 \pm 8.6$ | $-168.7 \pm 7.1$ | No (0.51) |
| 8. | K22R | $-581.6 \pm 29.2$ | $-589.9 \pm 26.6$ | Yes (<0.001) | $-131.6 \pm 6.8$ | $-134.0 \pm 6.0$ | Yes (<0.001) |
| 9. | RR | $-731.3 \pm 33.7$ | $-710.1 \pm 36.7$ | Yes (<0.001) | $-168.0 \pm 7.9$ | $-164.2 \pm 8.3$ | Yes (<0.001) |
| 10. | K22H | $-470.0 \pm 31.7$ | $-490.7 \pm 28.3$ | Yes (<0.001) | $-105.9 \pm 7.2$ | $-111.3 \pm 6.5$ | Yes (<0.001) |
| 11. | K22E | $-486.9 \pm 36.9$ | $-495.7 \pm 32.6$ | Yes (<0.001) | $-108.8 \pm 8.6$ | $-111.4 \pm 7.9$ | Yes (<0.001) |
| 12. | K22G | $-454.6 \pm 34.4$ | $-484.5 \pm 23.5$ | Yes (<0.001) | $-102.3 \pm 7.8$ | $-108.8 \pm 5.5$ | Yes (<0.001) |
| 13. | FLQ | $-481.4 \pm 23.4$ | $-476.2 \pm 22.4$ | Yes (<0.001) | $-108.2 \pm 5.4$ | $-106.6 \pm 5.1$ | Yes (<0.001) |

In order to understand why simulations with the two parameter sets lead to the above observed differences, we compare the polar component of solvation free energy (ΔG_sol,pol_). Again, we find only small differences in the mean values of ΔG_sol,pol_, but much larger ones in the standard deviations (Table 3 and Table S2). Similar behavior is seen for the non-polar component of solvation free energy (Table 3 and Table S2), and in general, arises the disparity in the ΔG_sol_ values from the additive effect of the smaller differences in the two components of ΔG_sol_. Only in a few cases, the differences arise from either the polar component (K22H, N20E, K22E, *etc.*) or the non-polar component (N20F) of solvation free energy.

**Table 3.**
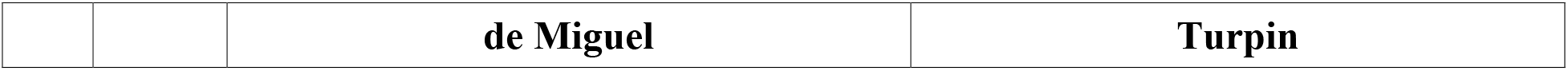
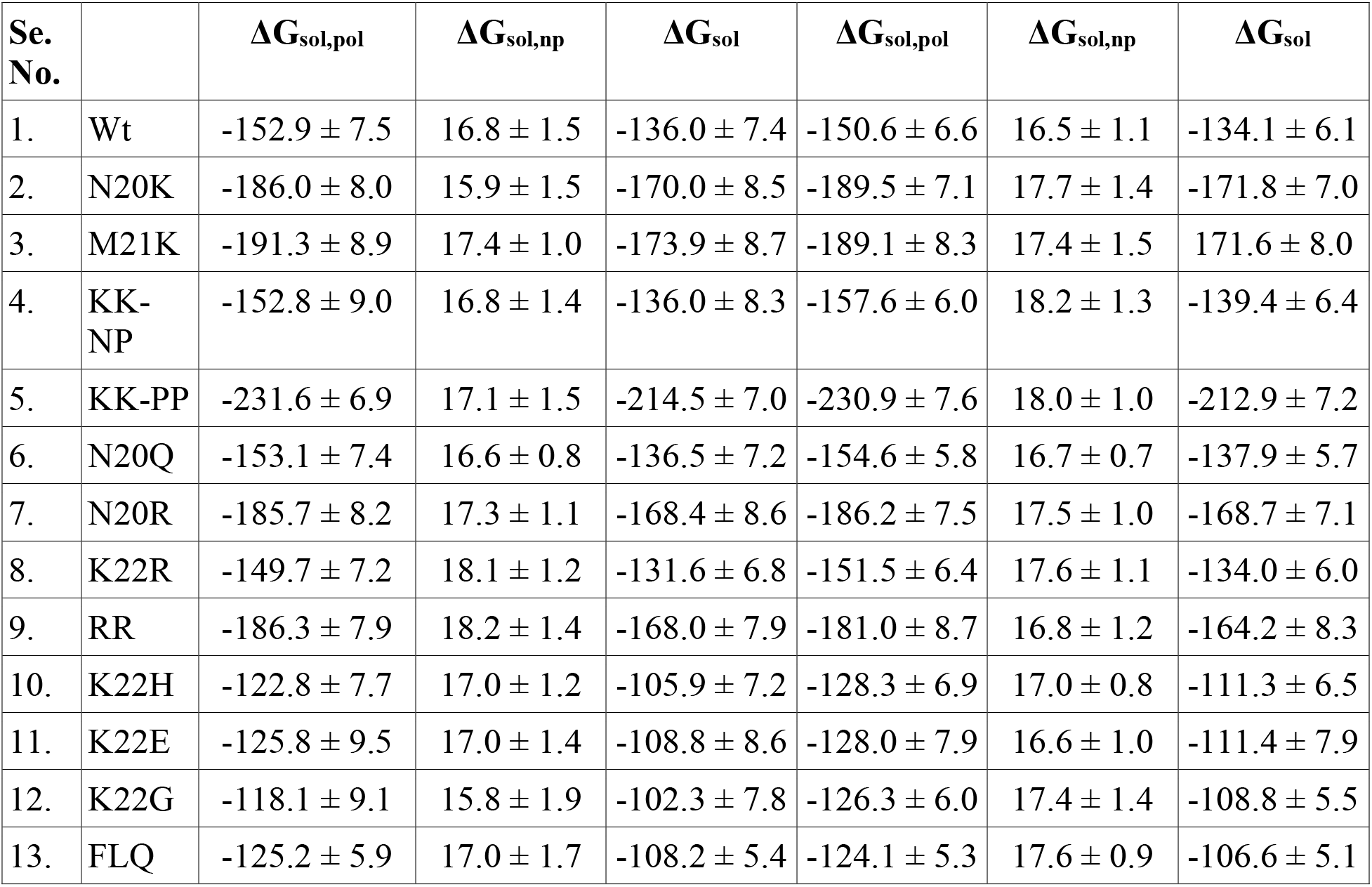
Calculated components of solvation free energy (ΔG_sol_) for nisin and mutants using parameters of de Miguel *et. al* and Turpin *et. al* using PBSA.

Since the significant differences in the calculated solubilities arise from a complex interplay between electrostatic and non-electrostatic interactions between peptide and solvent, it is not possible to decide *a priori* which of the two parameter sets is more appropriate. Experimentally, M21K, and N20K has been shown to be five-fold and three-fold more soluble than the wild-type, respectively. Hence, we have used tukey’s test to compare the relative order of solvation free energies (as calculated from simulations with the two parameter sets) with the experimental results. While with both parameters set, the mutants led to different solvation free energies than for the wild type, the Turpin *et. al* parameters could not differentiate between the two mutant forms. Only the de Miguel *et. al* parameters correctly predicted the order of the solubility of the two mutants (Figure 3 and Figure S2). At the same time, the double mutant N20K-M21K (de-protonated form) is experimentally known to be less soluble than the wild-type, but the Miguel *et. al* parameters could not distinguish mutant and wild type, and the Turpin *et. al* parameters even predicted higher solubility for the mutant. On the other hand, the solubility of the M21G and N20Q mutant could only be differentiated using Turpin *et. al* parameters while in case of the K22H mutant (slightly lower soluble than wild-type), both parameter sets appear to be equally reliable in predicting the order of solvation free energy (Figure 3 and Figure S2).

**Figure 3.**
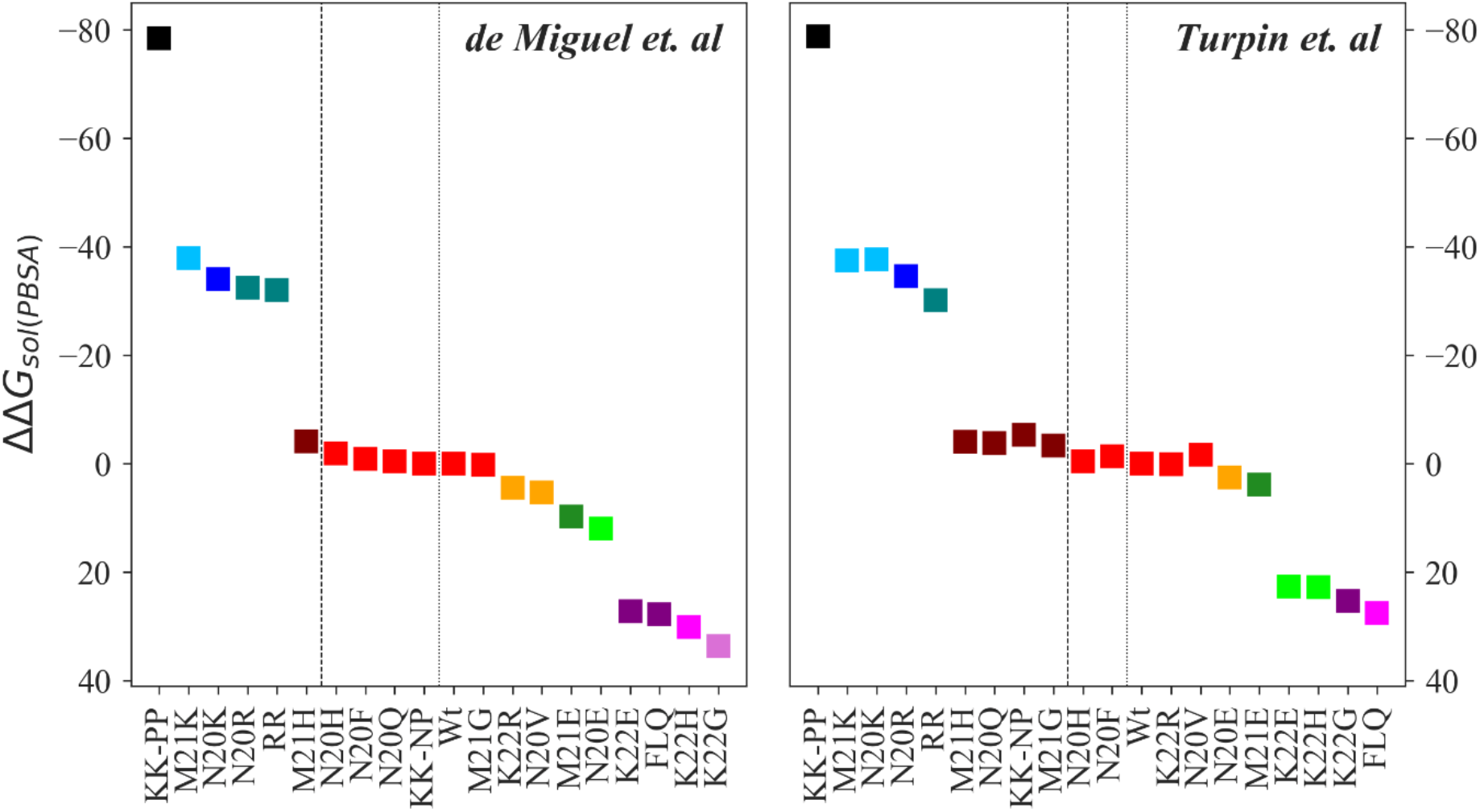
Effect of mutation on the solubility of Nisin estimated using PBSA. The figure shows the difference between the solvation free energies of mutant and wild-type nisin (ΔΔG_sol_ = ΔG_mutant_ ΔG_Wt_). Nisin and mutants are ranked in decreasing order of solubility. A different color is used for each rank, and a matching color is assigned to the mutants with the same rank. Statistical significance is determined by using multiple comparison Tukey’s test at α=0.05.

Hence, while the two force-fields allow one in most cases to predict qualitatively the relative solubility of mutants, they do differ significantly in the predicted solvation free energies, and the Turpin *et. al* parameters seems to favor more extended configurations than the de Miguel *et. al* parameters. We believe that the discrepancies in the calculated solvation free energy is a synergetic effect of the collapse of the polypeptide chain and dissimilarities in the first solvation shell resulting likely from the different partial atomic charges in the two parameter sets for the amide group and the carbonyl group of the dehydroamino acids (also the Cα atom). We have analyzed the collapse of the polypeptide chain and the behavior of water molecules by measuring the solvent-accessible surface area (SASA), number of peptide-water hydrogen bonds, number of water molecules in the first solvation shell (defined by a cut-off distance of 3.5 Å from the peptide surface), and the mean deviation of the water number fluctuation function (N_w_(t)), shown in Figure 4. The N_w_(t) is defined as

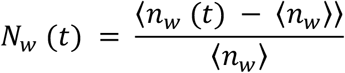

where, n_w_ (t) represents the number of water molecules in the first hydration shell at time ‘t’ and n_w_ is the average number of hydration water.

**Figure 4.**
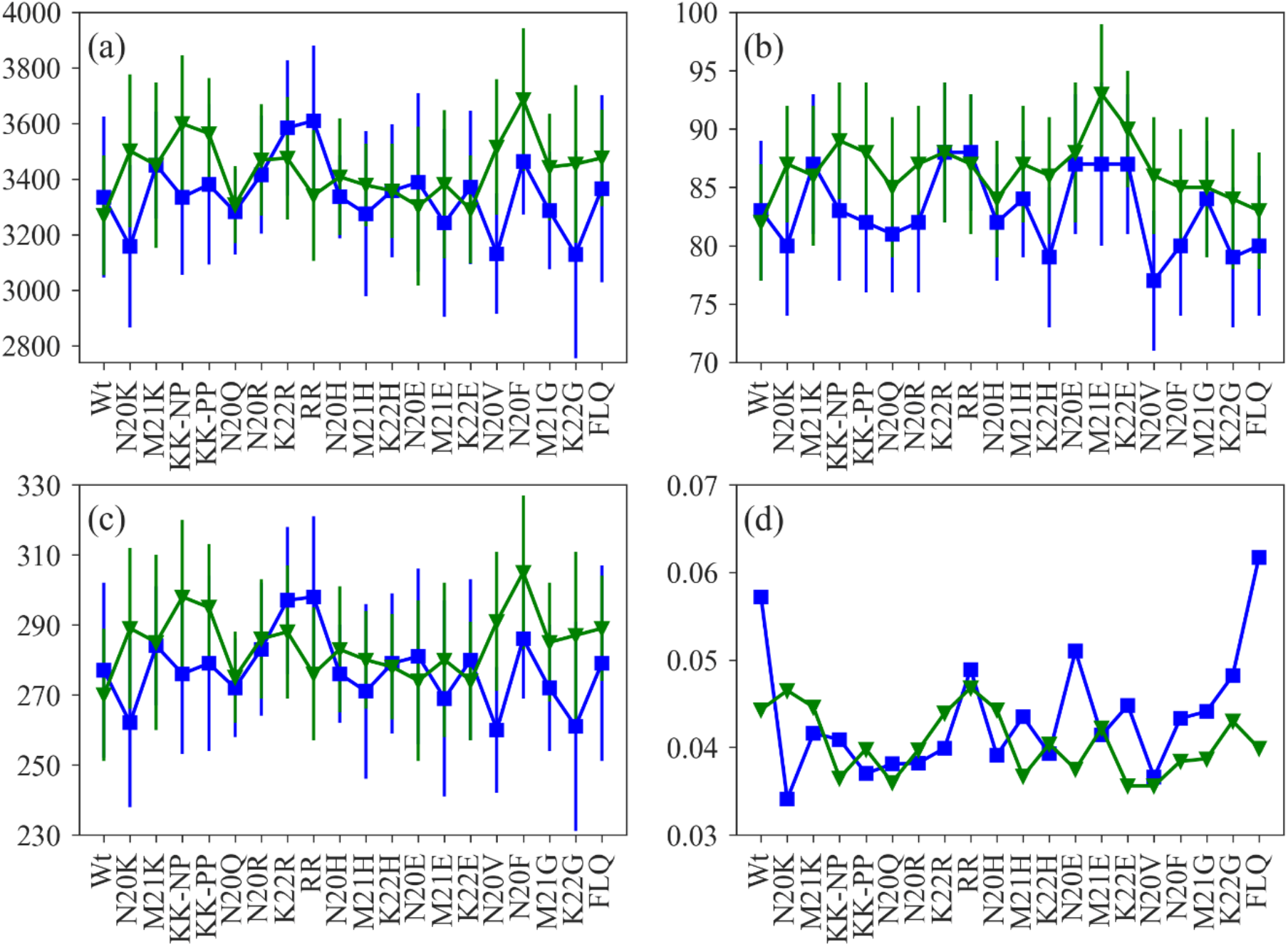
Collapse of the polypeptide chain and behavior of water molecules in the first hydration shell of nisin and mutants as a function of SASA (a), peptide-water hydrogen bond (b), number of water molecules in the first solvation shell (c) and mean deviation of water number fluctuation function (d). Color scheme: de Miguel *et. al* in blue and Turpin *et. al* in green.

Figure 4(a) shows that with the exception of K22R, RR and N20E, the solvent-accessible surface area (SASA) is larger for both wild type and mutants when using Turpin *et. al* parameters. The measured numbers of peptide-water hydrogen bonds in Figure 4(b) also indicate that for this parameter set nisin remains in an extended conformation. With the exception of K22R, RR and again N20E, the number of water molecules in the first hydration shell is higher when using the parameters of Turpin *et. al* (Figure 4(c)), although the mean deviation of the water number fluctuation function is lower (Figure 4(d)). If nisin, when using Turpin *et. al* parameters, mostly remains in an extended conformation, it is expected to have more water molecules in the first solvation shell. As in the Turpin *et. al* parameter set, the partial charges for the amide group of both dehydroamino acids take values of −0.67 (N) and 0.36 (H) have a larger magnitude than in the de Miguel *et. al* parameter set (−0.47 (N), 0.31 (H)), it is expected that the nisin variants form stronger hydrogen bonds to the hydrating water molecules. These results also corroborate with the mean deviation of water number fluctuation function, which is always low using Turpin *et. al* parameters (Figure 4(d)).

We conclude that the differences in both components of the solvation free energy likely arise from the differing behavior of the water molecules in the hydration layer of the peptide. As the hydration layer of the peptide using Turpin *et. al* parameter sets are more ordered and also the peptide-water association is stronger, it restricts the motion of the polypeptide chain, and therefore, we see less structural fluctuations using Turpin *et. al* parameters than de Miguel *et. al* parameters. However, while the two parameter sets sample different structural ensembles, they reproduce with a few exceptions *qualitatively* the experimentally measured solubility differences between wild-type and mutants, with no clear trend which of the two sets is more accurate.

We note that like for the amino group, the partial charges for the carbonyl group also differs for both the parameters sets (de Miguel *et. al*: 0.51 (C) and −0.51 (O); Turpin *et. al*: 0.635 (C) and −0.635 (O)). Varying these parameters and comparing calculated solvation free energies with the measured solubilities of the mutants, may allow to optimize the parameter sets. While our present investigation shows that the current two parameters sets agree with few exceptions in most cases with experimental solubility ranking. Further development of parameters either for the whole lantibiotics peptide or for the Dha and Dhb residues would be of significant value.^27–30^ While such a re-parametrization of the dehydroamino amino acids is beyond the scope of this paper, but our evaluation and comparison of the two parameter sets allows us already to list the issues that need to be addressed:

1. During force field parametrization, all isomeric forms need to be considered. For the parametrization of Dhb, de Miguel *et. al* considered only the Z-isomer, while Turpin *et. al* considered the E-isomer. Both forms are synthetized from the same precursor, threonine,^31–33^ and while in general the Z-isomer is more stable than the E isomer,^34^ the later is observed in several cases, alone or together with the Z-isomer.^35^
2. Both parameter sets do not use CMAP corrections. Since the torsion parameters strongly influence the molecular structures, it is necessary to provide the correct degree of rigidity as well as the flexibility to ensure and reproduce the significant conformational changes due to rotations about bonds. Hence, the torsion parameters used in these two parameter sets need to be refined.
3. There seems to be a disparity between the partial charges of the amide group and the carbonyl group of the dehydroamino acids (also the Cα atom) in both parameter sets, which affects the anchoring of the hydrating water molecules and therefore needs to be optimized.

### Effect of mutations on the solubility of nisin

While our above analysis has demonstrated significant shortcomings in the two parameter sets used to simulate nisin, it has shown at the same time that simulations with these sets allow one to reproduce *qualitatively* the experimentally observed differences in solubility between wild type and a few previously studied mutants. Hence, we conjecture that such simulations may already be sufficient to design new mutants that will be more soluble at physiological pH and, therefore, can serve as a basis for the future antibiotic design. Taking wild type nisin as our reference point, we want to understand the effect of various sidechain replacements on the solubility limit of the peptide in solution and use this knowledge to design new mutants with improved solubility. We estimate solubility again by the solvation free energy as approximated with the PBSA approach. We use both parameter sets (de Miguel and Turpin) in our analysis; however, the presented solvation free energy differences ΔΔG_sol_, between wild type and mutant, rely on the parameter set by Turpin *et. al*, as only this parameter set allowed to differentiate the N20Q and M21G mutants from the wild type. Furthermore, where the Turpin *et. al* parameter led to discrepancies with experimental solubility measurements, the computational results did not improve when using data from the simulations that relied on the de Miguel *et. al* parameters. We remark, however, that the ΔΔG_sol_ values are usually larger for positively charged mutations using Turpin *et. al* parameters, while the reverse is the case with the negatively charged mutations and mutation to histidine, *i.e.*, ΔΔG_sol_ values are higher when using de Miguel *et. al* parameters, and no clear trend in case of replacements to hydrophobic residues.

### Effect of mutations to positively charged residues

Introduction of a positively charged lysine at the positions 20, 21, and 20-21 results in a significant increase in the solubility of nisin (by a factor of 1.25-1.5 over the solubility of the wild-type, Figure 3). Especially the mutation to protonated lysine at position 20-21 (KK-PP) leads to a 1.5-fold increase in solubility (ΔΔG_sol_ = −78.8 kcal/mol), while replacement by the de-protonated lysine (KK-NP) resulted in little change (ΔΔG_sol_ = −5.3 kcal/mol, Figure 3). These results are consistent with earlier work in Refs.^8,9^ Replacements with positively charged arginine at positions 20 and 20-22 only moderately increased the solubility (~1.23 times, ΔΔG_sol_ N20R = −34.6 kcal/mol and ΔΔG_sol_ RR = −30.1 kcal/mol), and at position 22 did not enhance the solubility of nisin (ΔΔG_sol_ = 0.1 kcal/mol). Less effective are mutations to histidine (ΔΔG_sol_ ~ −4.0 – 22.8 kcal/mol, Figure 3), and the K22H mutant even decreases the solubility below that of the wild-type (ΔΔG_sol_ = 22.8 kcal/mol, Figure 3). Hence, mutations to a positively charged residue in the hinge region maximize the solubility of nisin, with lysine and arginine mutations to be the most effective ones. Of special importance here is our observation of the role of arginine mutations. Since the pK of titratable arginine group is high (~12.1), the mutants bearing arginine in the hinge region are less likely to be affected by the change in pH. We report three mutants (N20K^+^-M21K^+^, N20R-K22R, and N20R) with improved solubility compared to wild-type nisin.

### Effect of mutations to negatively charged residues

A mutation to glutamic acid in the hinge region has been investigated experimentally, showing a significant decrease or even loss in the production of nisin.^9^ It has been speculated that this loss is associated with an increase in steric hindrance in the hinge region. The assumption is that sidechain replacement to negatively charged amino acids will interfere with the thioether bridge formation (ring D), but further investigation is needed to determine the stages of biosynthesis that are blocked by such mutations. Nevertheless, even if such mutants with negatively charged side chains in the hinge region could be synthesized, a mutation to glutamic acid in the hinge region will likely not expand the solubility limit of nisin (K22E (ΔΔG_sol_ = 22.7 kcal/mol) < M21E (ΔΔG_sol_ = 3.9 kcal/mol) < N20E (ΔΔG_sol_ = 2.5 kcal/mol) < Wt), presumably due to the decrease in the net charge of the peptide. In particular, the solvation free energy difference shown in Figure 3 indicate that a mutation to glutamic acid at position 22 will decrease the solubility of nisin by about 20 %. Further, mutation to a negatively charged residue may also prevent the association of nisin and phospholipid bilayer as bacterial membranes carry a net negative surface charge, thereby decreasing the affinity of nisin. Hence, it appears that a positively charged residue such as arginine at position 22 is a better approach for expanding the solubility spectrum of nisin.

### Effect of mutations to uncharged polar and hydrophobic residues

While a mutation to an uncharged polar residue (N20Q) yields a slightly more soluble form of the peptide (ΔΔG_sol_ = −3.8 kcal/mol), mutations to a hydrophobic residue have little or no effect on nisin solubility (ΔΔG_sol_ ranges from −3.4 kcal/mol to −1.4 kcal/mol, Figure 3). An exception is a K22G mutant, which diminishes the solubility of nisin (ΔΔG_sol_ = 25.3 kcal/mol). A triple mutation (FLQ) designed to retain polarity at position 22, while increasing the hydrophobicity at two other positions, also only decreases the solubility of the peptide (ΔΔG_sol_ = 27.5 kcal/mol).

Hence, the hinge region (Asn-Met-Lys) in nisin controls not only the conformational flexibility needed for the antimicrobial activity, but also modulates the solubility of the peptide as summarized in the sketch of Figure 5. Especially effective are mutants introducing a positively charged residue into the hinge region. While introducing either a lysine or an arginine in the hinge region will increase the solubility, we believe that mutation to arginine will, in addition, increase nisin-membrane interaction, enhancing the antimicrobial activity of nisin. This is because the guanidium group of arginine binds more strongly to the phosphate groups of lipids than the amino group of lysine,^36^ and while bound to the lipid, the polarity on the guanidium group is reduced thereby increasing the capability to internalize membrane.^37,38^

**Figure 5.**
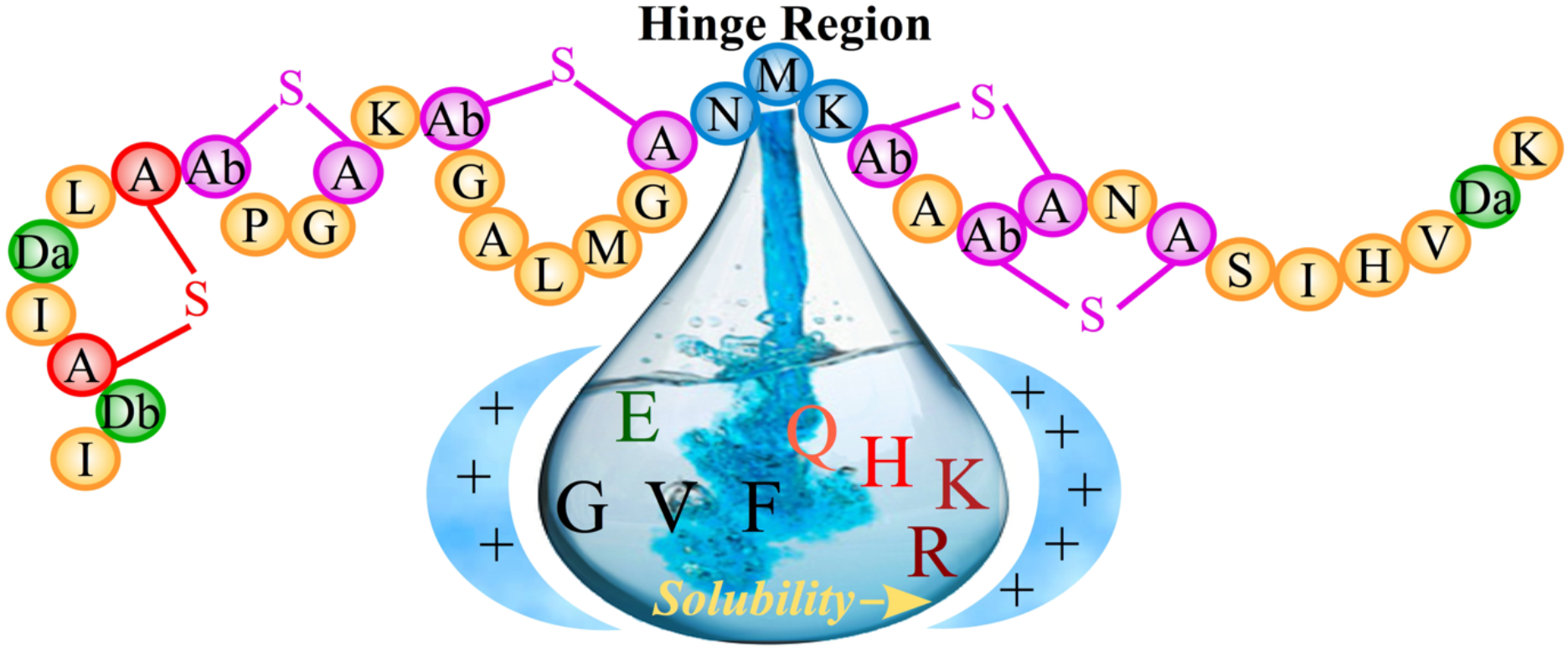
Schematic of effect of mutations in the hinge region on the solubility of nisin.

## Conclusion

Although nisin possesses excellent antimicrobial activity, its poor solubility and stability at physiological pH and temperature limit its use as a food preservative and even more as an antibiotic.^6,7^ Several efforts have been made in the past three decades to achieve a more soluble and antimicrobial active form of the peptide. While such a mutant would increase the range of conditions where nisin can be used as a food preservative, it would also open the door for possible uses of nisin as an antibiotic, as previous efforts have remained unsatisfactory. In this work, we use all-atom molecular dynamics simulations to understand the underlying mechanism for designing new mutants with improved solubility.

One of the challenges was the lack of standard CHARMM force field parameters for the two dehydroamino acids (dehydroalanine and dehydrobutyrine) and the thio-ether rings. Two sets of CHARMM compatible parameters have been proposed earlier that differ in the choice of the isomeric form of dehydrobutyrine, partial charges, and dihedral parameters. As in neither of the two studies were solubility calculations reported, we have first evaluated the relative merits of the two parameter sets for reproducing experimental solubility measurements for nisin. Estimating solubility by solvation free energies as approximated with the computationally cheap Poisson–Boltzmann and surface area continuum solvation (PBSA), we find that not only absolute values differ between the two parameter sets, but also are, in general, the corresponding standard deviations larger when using de Miguel *et. al* parameters. The choice of parameter set affects both components of the solvation free energy. We conjecture that the differences in the solvation free energy likely arise because the hydration layer of the peptide is more ordered and with stronger peptide-water association when using the Turpin *et. al* parameter set. While both parameter sets reproduce *qualitatively* the experimentally measured solubility differences between wild-type and mutants, our results do not allow us to select one parameter set over the other as being more accurate. Instead, they demonstrate the need for further improvements of the two parameter sets and pinpoint some of the shortcomings. Especially, we show the need to consider correct isomeric forms of dehydrobutyrine, dihedral parameters, and partial atomic charges for optimization of the force field parameters.

Since the existing parameter sets allow already for a qualitative assessment of solvation free energies, we used both sets in tandem to evaluate the change of solvation free energy and ranking of mutants. We observe that mutations to positively charged amino acid typically increases the solubility of the peptide, while the reverse is the case with the mutation to a negatively charged amino acid. Mutation to hydrophobic amino acids does not change systematically the solvation free energy, and mutations such as K22H, K22E, K22G, and N20F-M21L-K22Q that decrease the net charge also decrease the solubility. The effect of the various mutations is summarized in the sketch of Figure 5. Of special interest for possible applications are the new mutants N20R-K22R and N20R, as the high pKa (~12.1) ensures that arginine remains protonated under acidic, physiological, and alkaline pH without sacrificing the binding affinity of nisin towards the membrane. Hence, these mutants promise to extend the solubility of nisin over a broad range of pH values, therefore, broaden its use as a food preservative or potential antibiotics.

In conclusion, our study demonstrates that minor differences in force field parameters of even a few amino acids (Dha and Dhb, in case of nisin) can change dramatically the solubility of a peptide. This points to the need for a careful calibration of the force field parameters of the two dehydroamino acids not found in the standard CHARMM parameter set but common in antimicrobial peptides. Nevertheless, existing parameters allow already a *qualitative assessment of solvation free energy differences*, allowing us to propose new mutants, such as N20R-K22R and N20R, with potential applications as food preservative or antibiotics. In that, our study provides a basis for future antibiotic design.

## Supporting Information

The following Supporting Information is available free of charge at https://pubs.acs.org:

- Supplementary Tables: Calculated p-values of statistical test (Welch’s t-test) for solvation free energy (GBSA and PBSA) of Nisin and its mutant forms, Table S1; Calculated components of solvation free energy (ΔG_sol_) for nisin and mutants, Table S2.
- Supplementary Figures: Phi-psi (ϕ/ψ) backbone dihedral angle distribution of Dehydroalanine and Dehydrobutyrine, Figure S1; Effect of mutation on the solubility of Nisin estimated using GBSA, Figure S2.

## Acknowledgment

The simulations in this work were done using the SCHOONER cluster of the University of Oklahoma, and XSEDE resources allocated under grant MCB160005 (National Science Foundation). We acknowledge financial support from the National Institutes of Health under research grant GM120578.

## Table of Content Graphics

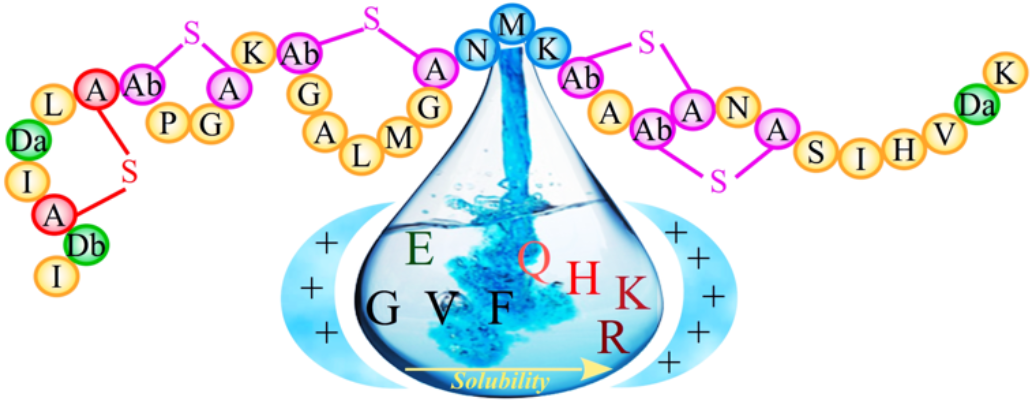

