## Supplementary information for "Altering the solubility of the antibiotic candidate Nisin – a computational study"

**Supplementary Tables**

**Table S1.** Calculated p-values of statistical test (Welch's t-test) for solvation free energy (GBSA and PBSA) of Nisin and its mutant forms using parameters of de Miguel *et. al* and Turpin *et. al.*

| Se. No. | | $\Delta G_{sol}$ (GBSA) | | | $\Delta G_{sol}$ (PBSA) ($\varepsilon_{in} = 4$) | | |
|---|---|---|---|---|---|---|---|
| | | **Miguel** | **Turpin** | **Significantly different? (p-value)** | **Miguel** | **Turpin** | **Significantly different? (p-value)** |
| 1. | WT | -594.3 ± 31.5 | -584.0 ± 26.8 | Yes (<0.001) | -136.0 ± 7.4 | -134.1 ± 6.1 | Yes (<0.001) |
| 2. | N20K | -732.2 ± 34.4 | -743.6 ± 29.7 | Yes (<0.001) | -170.0 ± 8.5 | -171.8 ± 7.0 | Yes (<0.001) |
| 3. | M21K | -752.3 ± 36.8 | -739.9 ± 34.6 | Yes (<0.001) | -173.9 ± 8.7 | -171.6 ± 8.0 | Yes (<0.001) |
| 4. | KK-NP | -592.9 ± 36.6 | -610.0 ± 25.7 | Yes (<0.001) | -136.0 ± 8.3 | -139.4 ± 6.4 | Yes (<0.001) |
| 5. | KK-PP | -917.1 ± 29.3 | -914.2 ± 31.9 | No (0.101) | -214.5 ± 7.0 | -212.9 ± 7.2 | Yes (<0.001) |
| 6. | N20Q | -595.6 ± 31.3 | -603.0 ± 24.4 | Yes (<0.001) | -136.5 ± 7.2 | -137.9 ± 5.7 | Yes (<0.001) |
| 7. | N20R | -731.4 ± 36.3 | -729.8 ± 31.1 | No (0.412) | -168.4 ± 8.6 | -168.7 ± 7.1 | No (0.51) |
| 8. | K22R | -581.6 ± 29.2 | -589.9 ± 26.6 | Yes (<0.001) | -131.6 ± 6.8 | -134.0 ± 6.0 | Yes (<0.001) |
| 9. | RR | -731.3 ± 33.7 | -710.1 ± 36.7 | Yes (<0.001) | -168.0 ± 7.9 | -164.2 ± 8.3 | Yes (<0.001) |
| 10. | N20H | -601.5 ± 25.0 | -588.9 ± 25.1 | Yes (<0.001) | -137.9 ± 5.9 | -134.6 ± 5.6 | Yes (<0.001) |
| 11. | M21H | -611.8 ± 27.1 | -602.9 ± 21.3 | Yes (<0.001) | -140.1 ± 6.3 | -138.1 ± 5.0 | Yes (<0.001) |
| 12. | K22H | -470.0 ± 31.7 | -490.7 ± 28.3 | Yes (<0.001) | -105.9 ± 7.2 | -111.3 ± 6.5 | Yes (<0.001) |
| 13. | N20E | -549.7 ± 58.3 | -576.5 ± 39.9 | Yes (<0.001) | -124.0 ± 13.9 | -131.6 ± 9.2 | Yes (<0.001) |
| 14. | M21E | -552.6 ± 47.2 | -571.9 ± 37.6 | Yes (<0.001) | -126.2 ± 10.9 | -130.2 ± 8.8 | Yes (<0.001) |
| 15. | K22E | -486.9 ± 36.9 | -495.7 ± 32.6 | Yes (<0.001) | -108.8 ± 8.6 | -111.4 ± 7.9 | Yes (<0.001) |
| 16. | N20V | -569.4 ± 42.2 | -595.6 ± 24.3 | Yes (<0.001) | -130.7 ± 9.5 | -135.8 ± 5.6 | Yes (<0.001) |
| 17. | N20F | -600.3 ± 29.3 | -596.8 ± 22.7 | Yes (0.021) | -136.9 ± 6.8 | -135.5 ± 5.2 | Yes (<0.001) |
| 18. | M21G | -593.1 ± 31.3 | -601.1 ± 20.6 | Yes (<0.001) | -135.8 ± 7.3 | -137.5 ± 4.9 | Yes (<0.001) |
| 19. | K22G | -454.6 ± 34.4 | -484.5 ± 23.5 | Yes (<0.001) | -102.3 ± 7.8 | -108.8 ± 5.5 | Yes (<0.001) |
| 20. | FLQ | -481.4 ± 23.4 | -476.2 ± 22.4 | Yes (<0.001) | -108.2 ± 5.4 | -106.6 ± 5.1 | Yes (<0.001) |

**Table S2.** Calculated components of solvation free energy ($\Delta G_{sol}$) for nisin and mutants using parameters of de Miguel *et. al* and Turpin *et. al* using PBSA.

| Se. No. | | Miguel | | | Turpin | | |
|---|---|---|---|---|---|---|---|
| | | $\Delta G_{sol,pol}$ | $\Delta G_{sol,np}$ | $\Delta G_{sol,pol}$ | $\Delta G_{sol,np}$ | $\Delta G_{sol,pol}$ | $\Delta G_{sol,np}$ |
| 1. | WT | -152.9 ± 7.5 | 16.8 ± 1.5 | -136.0 ± 7.4 | -150.6 ± 6.6 | 16.5 ± 1.1 | -134.1 ± 6.1 |
| 2. | N20K | -186.0 ± 8.0 | 15.9 ± 1.5 | -170.0 ± 8.5 | -189.5 ± 7.1 | 17.7 ± 1.4 | -171.8 ± 7.0 |
| 3. | M21K | -191.3 ± 8.9 | 17.4 ± 1.0 | -173.9 ± 8.7 | -189.1 ± 8.3 | 17.4 ± 1.5 | 171.6 ± 8.0 |
| 4. | KK-NP | -152.8 ± 9.0 | 16.8 ± 1.4 | -136.0 ± 8.3 | -157.6 ± 6.0 | 18.2 ± 1.3 | -139.4 ± 6.4 |
| 5. | KK-PP | -231.6 ± 6.9 | 17.1 ± 1.5 | -214.5 ± 7.0 | -230.9 ± 7.6 | 18.0 ± 1.0 | -212.9 ± 7.2 |
| 6. | N20Q | -153.1 ± 7.4 | 16.6 ± 0.8 | -136.5 ± 7.2 | -154.6 ± 5.8 | 16.7 ± 0.7 | -137.9 ± 5.7 |
| 7. | N20R | -185.7 ± 8.2 | 17.3 ± 1.1 | -168.4 ± 8.6 | -186.2 ± 7.5 | 17.5 ± 1.0 | -168.7 ± 7.1 |
| 8. | K22R | -149.7 ± 7.2 | 18.1 ± 1.2 | -131.6 ± 6.8 | -151.5 ± 6.4 | 17.6 ± 1.1 | -134.0 ± 6.0 |
| 9. | RR | -186.3 ± 7.9 | 18.2 ± 1.4 | -168.0 ± 7.9 | -181.0 ± 8.7 | 16.8 ± 1.2 | -164.2 ± 8.3 |
| 10. | N20H | -154.8 ± 5.9 | 16.9 ± 0.8 | -137.9 ± 5.9 | -151.8 ± 6.1 | 17.2 ± 1.1 | -134.6 ± 5.6 |
| 11. | M21H | -156.6 ± 6.6 | 16.5 ± 1.5 | -140.1 ± 6.3 | -155.1 ± 5.1 | 17.0 ± 0.7 | -138.1 ± 5.0 |
| 12. | K22H | -122.8 ± 7.7 | 17.0 ± 1.2 | -105.9 ± 7.2 | -128.3 ± 6.9 | 17.0 ± 0.8 | -111.3 ± 6.5 |
| 13. | N20E | -141.1 ± 15.1 | 17.1 ± 1.6 | -124.0 ± 13.9 | -148.3 ± 10.3 | 16.7 ± 1.5 | -131.6 ± 9.2 |
| 14. | M21E | -142.5 ± 12.1 | 16.4 ± 1.7 | -126.2 ± 10.9 | -147.3 ± 9.2 | 17.0 ± 1.3 | -130.2 ± 8.8 |
| 15. | K22E | -125.8 ± 9.5 | 17.0 ± 1.4 | -108.8 ± 8.6 | -128.0 ± 7.9 | 16.6 ± 1.0 | -111.4 ± 7.9 |
| 16. | N20V | -146.5 ± 9.9 | 15.8 ± 1.1 | -130.7 ± 9.5 | -153.6 ± 6.1 | 17.8 ± 1.2 | -135.8 ± 5.6 |
| 17. | N20F | -154.4 ± 7.1 | 17.5 ± 0.9 | -136.9 ± 6.8 | -154.2 ± 5.6 | 18.7 ± 1.3 | -135.5 ± 5.2 |
| 18. | M21G | -152.4 ± 7.6 | 16.6 ± 1.1 | -135.8 ± 7.3 | -155.0 ± 5.0 | 17.4 ± 1.0 | -137.5 ± 4.9 |
| 19. | K22G | -118.1 ± 9.1 | 15.8 ± 1.9 | -102.3 ± 7.8 | -126.3 ± 6.0 | 17.4 ± 1.4 | -108.8 ± 5.5 |
| 20. | FLQ | -125.2 ± 5.9 | 17.0 ± 1.7 | -108.2 ± 5.4 | -124.1 ± 5.3 | 17.6 ± 0.9 | -106.6 ± 5.1 |

### Supplementary Figures

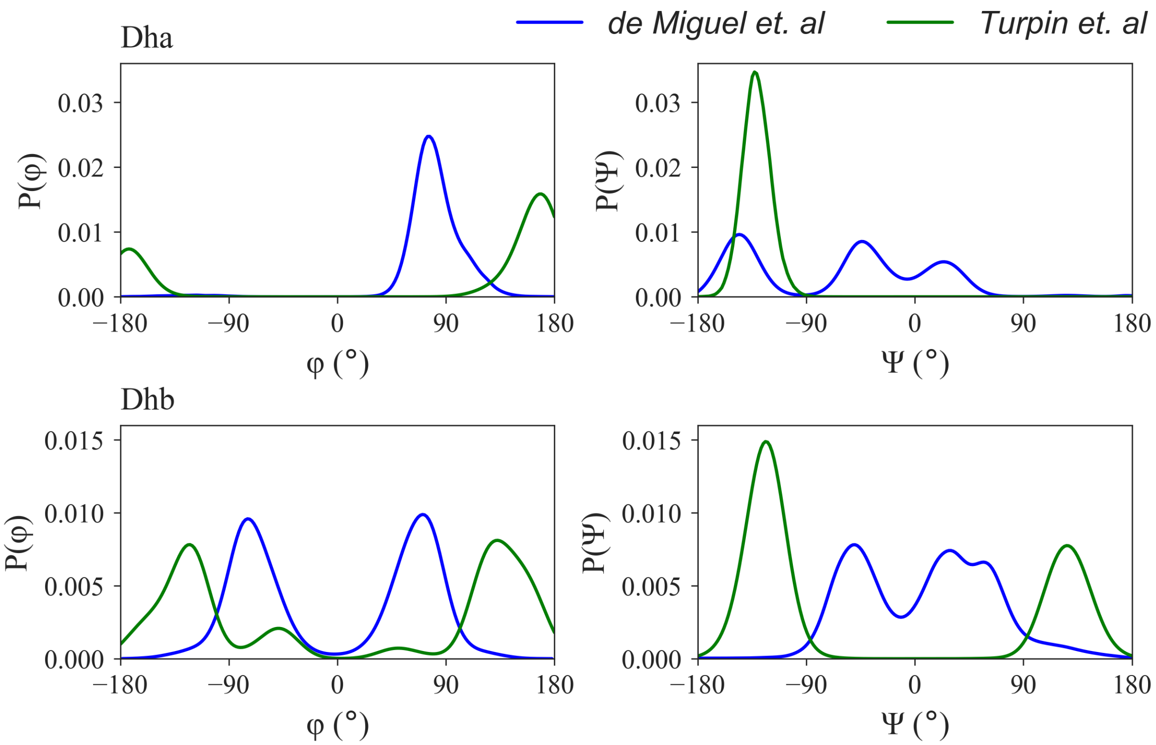


**Figure S1.** Phi-psi (ϕ/ψ) backbone dihedral angle distribution of Dehydroalanine and Dehydrobutyrine.

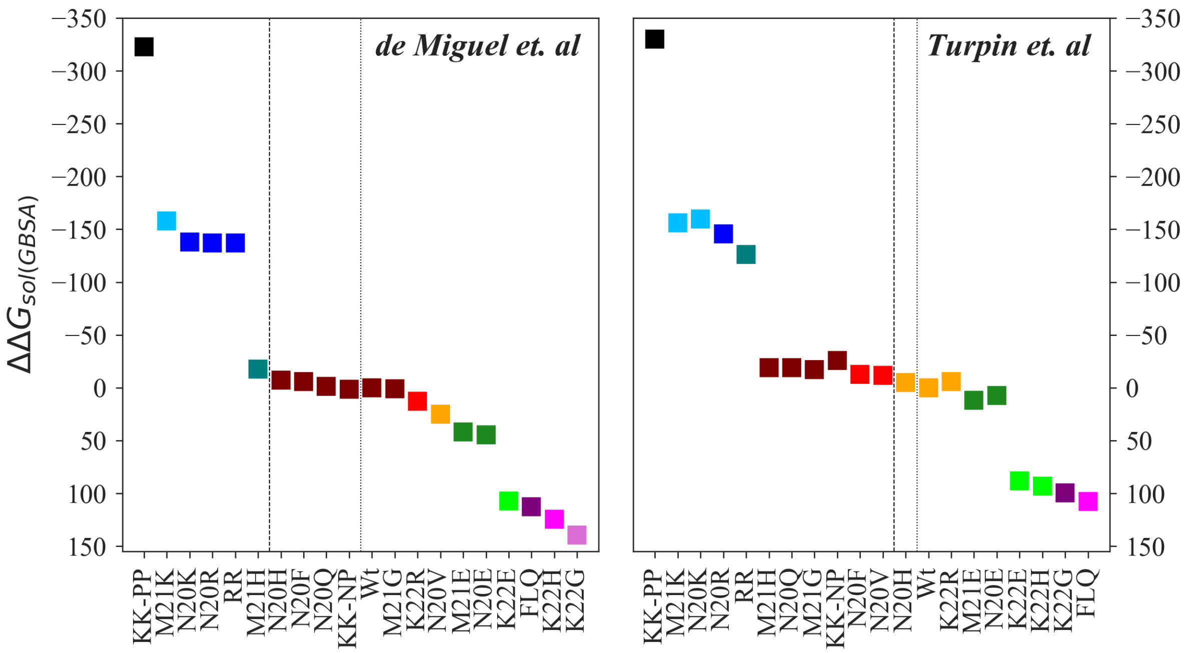


**Figure S2.** Effect of mutation on the solubility of Nisin estimated using GBSA. The figure shows the difference between the solvation free energies of mutant and wild-type nisin ($\Delta\Delta G_{sol} = \Delta G_{mutant} - \Delta G_{Wt}$). Nisin and mutants are ranked in decreasing order of solubility. A different color is used for each rank, and a matching color is assigned to the mutants with the same rank. Statistical significance is determined by using multiple comparison Tukey's test at $\alpha$=0.05.
